# SARS-CoV-2 and ORF3a: Non-Synonymous Mutations and Polyproline Regions

**DOI:** 10.1101/2020.03.27.012013

**Authors:** Elio Issa, Georgi Merhi, Balig Panossian, Tamara Salloum, Sima Tokajian

**Author notes:** Address correspondence to Sima T. Tokajian, PhD, Department of Natural Sciences, School of Arts and Sciences, Lebanese American University, Byblos, 36, Lebanon. E.I, G.M and B.P contributed equally to this work.

## Abstract

The effect of the rapid accumulation of non-synonymous mutations on the pathogenesis of SARS-CoV-2 is not yet known. To predict the impact of non-synonymous mutations and polyproline regions identified in ORF3a on the formation of B-cell epitopes and their role in evading the immune response, nucleotide and protein sequences of 537 available SARS-CoV-2 genomes were analyzed for the presence of non-synonymous mutations and polyproline regions. Mutations were correlated with changes in epitope formation. A total of 19 different non-synonymous amino acids substitutions were detected in ORF3a among 537 SARS-CoV-2 strains. G251V was the most common and identified in 9.9% (n=53) of the strains and was predicted to lead to the loss of a B-cell like epitope in ORF3a. Polyproline regions were detected in two strains (EPI_ISL_410486, France and EPI_ISL_407079, Finland) and affected epitopes formation. The accumulation of non-synonymous mutations and detected polyproline regions in ORF3a of SARS-CoV-2 could be driving the evasion of the host immune response thus favoring viral spread. Rapid mutations accumulating in ORF3a should be closely monitored throughout the COVID-19 pandemic.

**Importance:** At the surge of the COVID-19 pandemic and after three months of the identification of SARS-CoV-2 as the disease-causing pathogen, nucleic acid changes due to host-pathogen interactions are insightful into the evolution of this virus. In this paper, we have identified a set of non-synonymous mutations in ORF3a and predicted their impact on B-cell like epitope formation. The accumulation of non-synonymous mutations in ORF3a could be driving protein changes that mediate immune evasion and favoring viral spread.

## Introduction

The rapid spread of the coronavirus disease 2019 (COVID-19) caused by a novel coronavirus, named SARS-CoV-2 due to its symptoms similarity to those induced by the severe acute respiratory syndrome (SARS), is a major global concern (1). The epidemic started in late December 2019 in Wuhan, the capital of Central China’s Hubei Province and since then thousands of cases have been reported in more than 46 countries (https://www.who.int/emergencies/diseases/novel-coronavirus-2019/situation-reports/). Coronaviruses are enveloped non-segmented positive sense RNA viruses belonging to the family Coronaviridae and the order Nidovirales and are broadly distributed in humans and other mammals. The genome of SARS-CoV-2 showed 96.2% sequence similarity to a bat SARS-related coronavirus (SARSr-CoV; RaTG13) collected in Yunnan province, China (1) and 79% and 50% similarities to SARS-CoV and MERS-CoV, respectively (2). A transmission from wild-life animals (such as pangolins) to humans has been recently suggested (3).

With the immediate and continuous release of sequence data, monitoring the rapid evolution of the SARS-CoV-2 genome provides a strong lead towards predicting and potentially mitigating its global spread. ORF3a protein (Accession # YP_009724391.1) is a hypothetical protein showing a 72% sequence similarity to SARS3a protein in SARS-CoV. Here, we investigated the presence of diverse non-synonymous mutations in ORF3a and their effects on the predicted protein structure and its potential implication in the formation of epitopes. Moreover, polyproline regions (PPRs) were detected in two strains. We used this approach to follow and understand the impact of new emerging mutations in the pathogenesis and immune evasion of SARS-CoV-2.

## Results

### Micro-clonality within ORF3a

The clonal diversity of SARS-CoV-2 core genomes was highly similar in tree topology to the gene tree of ORF3a (Figure 1). Signature mutations within SARS-CoV-2 genomes cluster them into defined phylogenetic clades. Similarly, we observed micro-clonality within the ORF3a gene tree defined by highlighted non-synonymous mutations G251V (green) and Q57H (pink) that are found in conserved phylogenetic micro-clades representing sub-populations of mutants.

**Figure 1:**
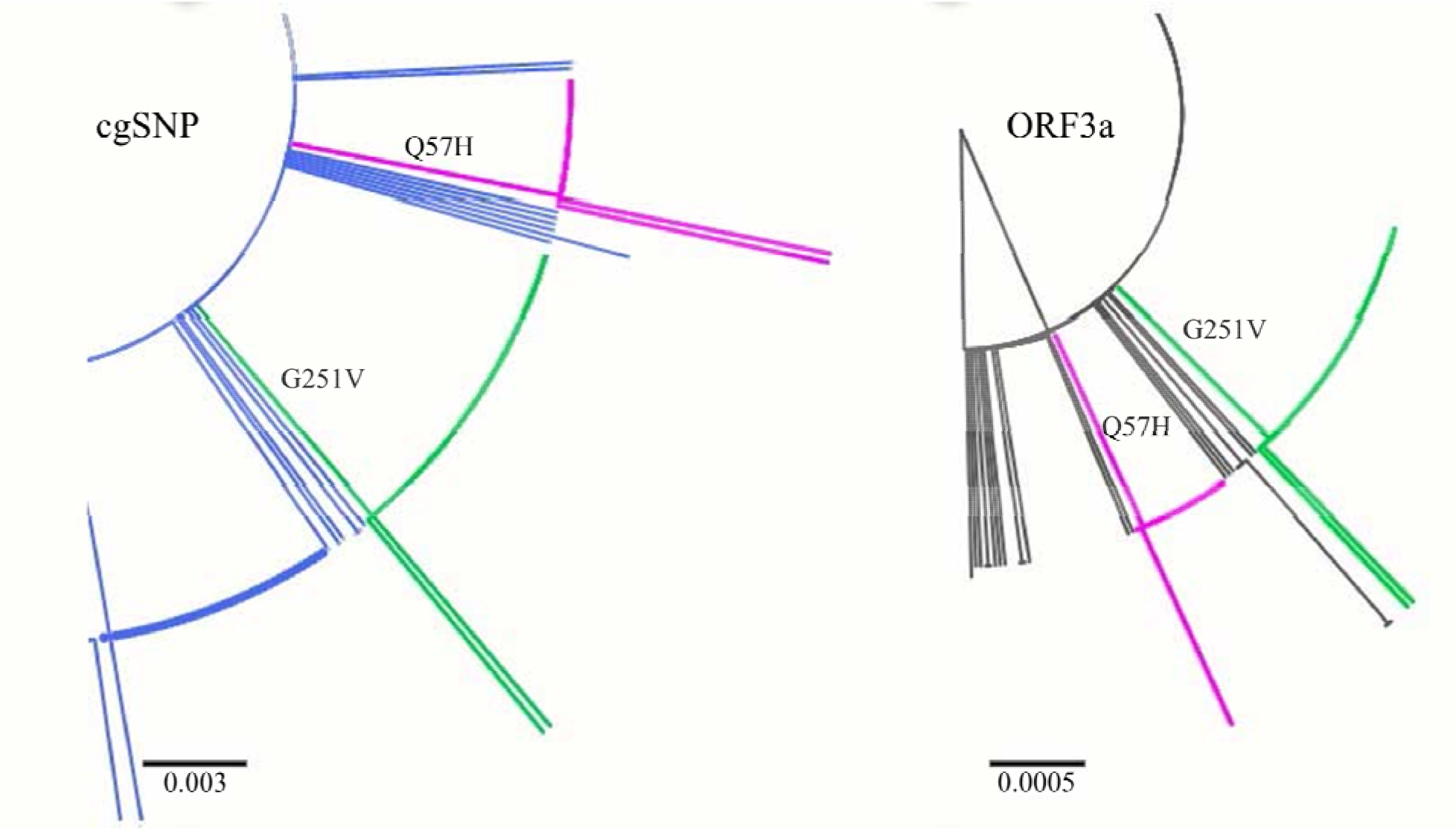
Phylogenetic trees of SARS-CoV-2 core genomes and ORF3a. Magnified maximum-likelihood phylogenetic trees of **(A)** SARS-CoV-2 genomes based on core genome SNP differences in all concatenated ORFs and **(B)** ORF3a gene tree highlighting G251V mutant clade in green and Q57H mutant clade in pink.

### Non-synonymous Mutations in ORF3a

ORF3a, encoding a hypothetical protein, showed a 97.82% sequence similarity (100% coverage) to a nonstructural protein NS3 of Bat coronavirus RaTG13 (Accession # QHR63301.1). Moreover, ORF3a has a pro-apoptosis inducing APA3_viroporin conserved domain, also found in SARS-CoV.

Sequence alignment of 537 ORF3a protein sequences revealed a total of 19 non-synonymous amino acids substitutions, of which 52.6% (n=10) had a predicted deleterious functional outcome and 47.4% (n=9) had a neutral functional outcome (Figure 2.A).

**Figure 2:**
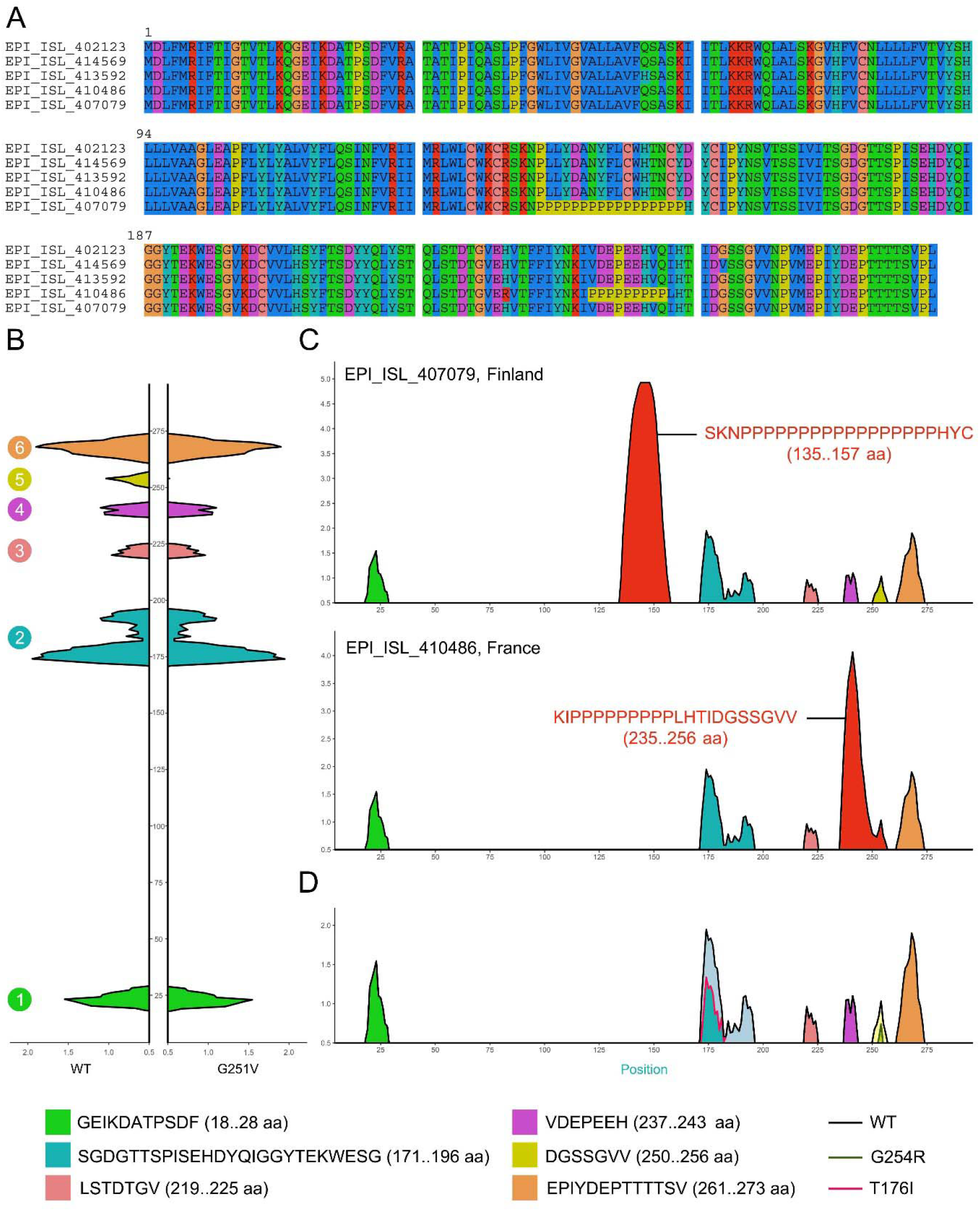
Mutations analysis of ORF3a. **(A)** Multiple sequence alignment between ORF3a protein of G251V (EPI_ISL_414569, Hong Kong), G254R (EPI_ISL_415627, USA), T176I (EPI_ISL_411950, Jiangsu), PPR-containing proteins (EPI_ISL_410486, France and EPI_ISL_407079, Finland) mutants compared to non-mutant (EPI_ISL_402123, Wuhan) **(B)** B-cell like epitopes of the non-mutated ORF3a protein (left) and G251V mutant (right). Only values above the threshold (0.5) are included. The mutation lead to the loss of one B cell epitope. **(C)** B-cell like epitopes of PPR-containing isolates. Additional epitopes are indicated in red. **(D)** B-cell like epitopes of T176I and G254R mutants with decreased intensity as compared to non-mutant.

G251V was the most frequently detected substitution found in 9.9% (n=53) of the strains followed by Q57H found in 3.9% (n=21) of the strains. Both G251V and Q57H were predicted to be deleterious (Table 1).

**Table 1.**
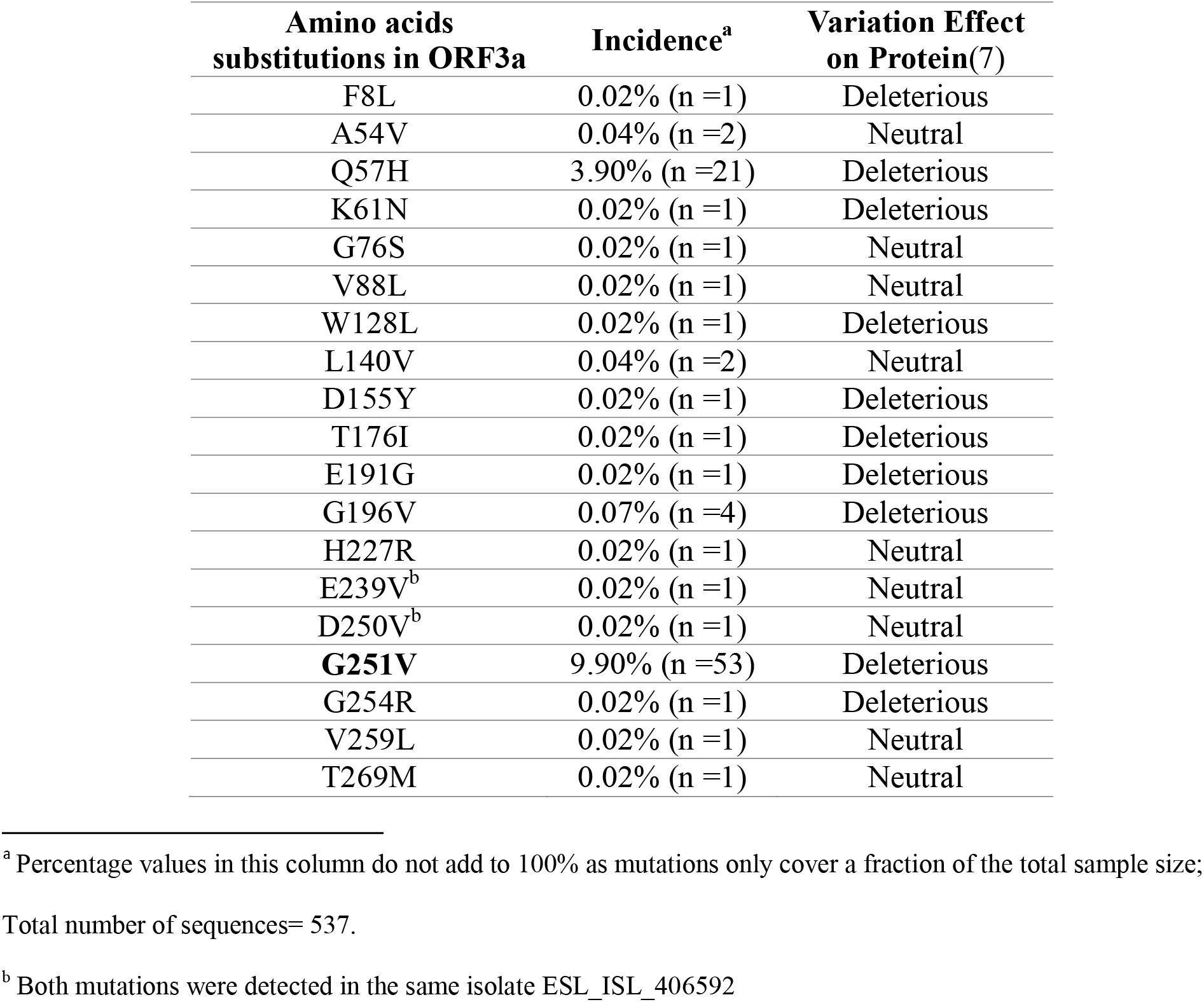
List of 19 non-synonymous amino acids substitutions in ORF3a among 537 strains. The G251V substitution is shown in bold.

### G251V linked to an Epitope Loss

The G251V mutations were further investigated. Motif scanning demonstrated that G251V resulted in the loss of a phosphatidylinositol-specific phospholipase X-box domain (PIPI_CX_DOMAIN; 203-275 aa). The G251V substitution created serine protease cleavage site. IEDB analysis revealed the presence of six putative epitopes in the non-mutant ORF3a compared to five epitopes in the mutant ORF3a (Figure 2.B). The G251V substitution in ORF3a was linked to the loss of a putative epitope the impact of which on viral spread and pathogenesis requires further experimental studies. Other T176I and G254R substitutions resulted in a decreased intensity of epitopes number two (blue) and epitope number five (yellow) (Figure 2.D).

### Detection of PPRs

Notably, we detected PPRs in two SARS-CoV-2 genomes (EPI_ISL_407079, Finland and EPI_ISL_410486, France). PPRs resulted in the joining of epitopes number four (purple) and five (yellow) into one larger epitope(red) of 22 amino acids in size (start:235; end:256; sequence: KIPPPPPPPPPLHTIDGSSGVV) in EPI_ISL_410486 (France) and led the appearance of a new epitope 23 amino acids in size (start 135; end 157; sequence: SKNPPPPPPPPPPPPPPPPHYC) in EPI_ISL_407079 (Finland) (Figure 2.C). Blastn search of a 23 bp DNA stretch from non-mutant strains showed a 100% identity to RaTG13 (Accession # MN996532.1).

## Discussion

These combined results suggest that the non-synonymous G251V mutation introduced into ORF3a protein in SARS-CoV-2 could be linked to immune evasion and thus viral spread and pathogenesis. ORF3a is a transmembrane protein that localizes to the plasma membrane especially in the ER-Golgi region and activates the PKR-like ER kinase (PERK) signaling pathway which protects viral proteins against ER-associated degradation. The activation of this pathway leads to apoptosis(11). A pro-apoptosis inducing APA3_viroporin conserved domain detected in ORF3a of SARS-CoV-2 is also found in SARS-CoV 3A protein (11).

The G251V was detected in ORF3a in 9.9% of the strains (n=53). G251V led to the loss a B cell-like epitope and a PIPLC_X_DOMAIN the eukaryotic homologue of which is involved in signal transduction processes (8). The accumulation of non-synonymous mutations could be driven by the humoral immunity as reported previously in the mucin-like domain of the Ebola virus glycoprotein (12).

Of paramount importance is the emergence of PPRs in ORF3a detected in two of the SARS-CoV-2 sequenced genomes (in EPI_ISL_410486, France and EPI_ISL_407079, Finland). PPRs are an open field for recombination that viruses use to adapt based on selective pressure (13). PPRs were previously shown to be indispensable for the activity of the Coxsackievirus B 3A protein which blocks ER-to-Golgi transport affecting protein synthesis (14). Studies on Hepatitis E virus also highlighted the role of PPRs in host-range adaptation and viral replication (15).

In conclusion, our study reveals and for the first time a common non-synonymous G251V substitution and PPRs in ORF3a which could be respectively linked to the loss of a putative epitope and viral spread and pathogenesis.

## Materials and Methods

### Pan-genome analysis

A total of 537 SARS-CoV-2 complete genomes with high quality sequencing downloaded from GISAID were utilized for genome and ORF3a alignments.

All sequences were uniformly annotated using Prokka v 1.1.3 (4). The annotated Genbank files were edited to have more concise locus tag identifiers. The Genbank annotations of the genomes were used as input in the PanX (5) pipeline for pan genome analysis. A core genome threshold of 0.99, MCL inflation parameter of 1.5, and a modified core diversity cutoff for branch lengths above 0.001 were used alongside the default parameters.

### Protein Structure prediction

Sequences were aligned using MUSCLE v3.8.31 (6). PROVEAN was used to predict the functional effects of amino acid substitutions (7). ExPASy and PROSPER were used for motif scanning and protease site prediction, respectively (8, 9). The Immune epitope database analysis resource (IEDB-AR) was used for epitopes prediction using a 0.5 threshold and default settings (10).

## Acknowledgements

We thankfully acknowledge the authors, generating and submitting laboratories of the sequences from GISAID’s EpiCoV™ database. We also acknowledge the authors of all Coronaviridae genome sequences deposited in GenBank. This study does not claim ownership of these sequences, which were used within the analysis workflow to further our understanding of the on-going pandemic of SARS-CoV-2 and the underlying molecular changes that govern the virus’ transmission and infectivity patterns. The authors wish to declare that they do not have any conflict of interests.

## Author Contributions

*Concept and design:* S.T. *Acquisition, analysis, or interpretation of data:* All authors. *Drafting of the manuscript:* All authors. *Critical revision of the manuscript for important intellectual content:* Tokajian. *Administrative, technical, or material support:* E.I, B.P, G.M. *Supervision:* S.T.

## References

1. Zhou P, Yang X-L, Wang X-G, Hu B, Zhang L, Zhang W, Si H-R, Zhu Y, Li B, Huang C-L, Chen H-D, Chen J, Luo Y, Guo H, Jiang R-D, Liu M-Q, Chen Y, Shen X-R, Wang X, Zheng X-S, Zhao K, Chen Q-J, Deng F, Liu L-L, Yan B, Zhan F-X, Wang Y-Y, Xiao G-F, Shi Z-L. 2020. A pneumonia outbreak associated with a new coronavirus of probable bat origin. Nature 1–4.

2. Gralinski LE, Menachery VD. 2020. Return of the Coronavirus: 2019-nCoV. 2. Viruses 12:135.

3. Andersen KG, Rambaut A, Lipkin WI, Holmes EC, Garry RF. 2020. The proximal origin of SARS-CoV-2. Nat Med 1–3.

4. Seemann T. 2014. Prokka: rapid prokaryotic genome annotation. Bioinforma Oxf Engl 30:2068–2069.

5. Ding W, Baumdicker F, Neher RA. 2018. panX: pan-genome analysis and exploration. Nucleic Acids Res 46:e5.

6. Edgar RC. 2004. MUSCLE: multiple sequence alignment with high accuracy and high throughput. Nucleic Acids Res 32:1792–1797.

7. Choi Y, Chan AP. 2015. PROVEAN web server: a tool to predict the functional effect of amino acid substitutions and indels. Bioinformatics 31:2745–2747.

8. Artimo P, Jonnalagedda M, Arnold K, Baratin D, Csardi G, de Castro E, Duvaud S, Flegel V, Fortier A, Gasteiger E, Grosdidier A, Hernandez C, Ioannidis V, Kuznetsov D, Liechti R, Moretti S, Mostaguir K, Redaschi N, Rossier G, Xenarios I, Stockinger H. 2012. ExPASy: SIB bioinformatics resource portal. Nucleic Acids Res 40:W597–W603.

9. Song J, Li F, Leier A, Marquez-Lago TT, Akutsu T, Haffari G, Chou K-C, Webb GI, Pike RN. 2018. PROSPERous: high-throughput prediction of substrate cleavage sites for 90 proteases with improved accuracy. Bioinformatics 34:684–687.

10. Zhang Q, Wang P, Kim Y, Haste-Andersen P, Beaver J, Bourne PE, Bui H-H, Buus S, Frankild S, Greenbaum J, Lund O, Lundegaard C, Nielsen M, Ponomarenko J, Sette A, Zhu Z, Peters B. 2008. Immune epitope database analysis resource (IEDB-AR). Nucleic Acids Res 36:W513–518.

11. Minakshi R, Padhan K, Rani M, Khan N, Ahmad F, Jameel S. 2009. The SARS Coronavirus 3a protein causes endoplasmic reticulum stress and induces ligand-independent downregulation of the type 1 interferon receptor. PloS One 4:e8342.

12. Park DJ, Dudas G, Wohl S, Goba A, Whitmer SLM, Andersen KG, Sealfon RS, Ladner JT, Kugelman JR, Matranga CB, Winnicki SM, Qu J, Gire SK, Gladden-Young A, Jalloh S, Nosamiefan D, Yozwiak NL, Moses LM, Jiang P-P, Lin AE, Schaffner SF, Bird B, Towner J, Mamoh M, Gbakie M, Kanneh L, Kargbo D, Massally JLB, Kamara FK, Konuwa E, Sellu J, Jalloh AA, Mustapha I, Foday M, Yillah M, Erickson BR, Sealy T, Blau D, Paddock C, Brault A, Amman B, Basile J, Bearden S, Belser J, Bergeron E, Campbell S, Chakrabarti A, Dodd K, Flint M, Gibbons A, Goodman C, Klena J, McMullan L, Morgan L, Russell B, Salzer J, Sanchez A, Wang D, Jungreis I, Tomkins-Tinch C, Kislyuk A, Lin MF, Chapman S, MacInnis B, Matthews A, Bochicchio J, Hensley LE, Kuhn JH, Nusbaum C, Schieffelin JS, Birren BW, Forget M, Nichol ST, Palacios GF, Ndiaye D, Happi C, Gevao SM, Vandi MA, Kargbo B, Holmes EC, Bedford T, Gnirke A, Ströher U, Rambaut A, Garry RF, Sabeti PC. 2015. Ebola Virus Epidemiology, Transmission, and Evolution during Seven Months in Sierra Leone. Cell 161:1516–1526.

13. Lhomme S, Abravanel F, Dubois M, Sandres-Saune K, Mansuy J-M, Rostaing L, Kamar N, Izopet J. 2014. Characterization of the polyproline region of the hepatitis E virus in immunocompromised patients. J Virol 88:12017–12025.

14. Wessels E, Duijsings D, Notebaart RA, Melchers WJG, Kuppeveld FJM van. 2005. A Proline-Rich Region in the Coxsackievirus 3A Protein Is Required for the Protein To Inhibit Endoplasmic Reticulum-to-Golgi Transport. J Virol 79:5163–5173.

15. Purdy MA, Lara J, Khudyakov YE. 2012. The Hepatitis E Virus Polyproline Region Is Involved in Viral Adaptation. PLOS ONE 7:e35974.

